# BiosInt: Biosensor-based smart design of pathway dynamic regulation for industrial biomanufacturing

**DOI:** 10.1101/2025.11.17.688676

**Authors:** Hèctor Martín Lázaro, Irene Otero-Muras, Pablo Carbonell

## Abstract

Industrial-scale production of bio-based chemicals in the circular green bioeconomy still faces inefficiencies arising from scaling up challenges. Biosensor mediated control of metabolic pathways has been proposed as a strategy to improve the performance of those bioprocesses. By linking industrial biosynthetic pathways to measurable metabolites through biochemical transformations, a highly interconnected regulatory design space is unveiled, offering new opportunities for pathway dynamic control. Yet, a systematic approach for selecting and implementing genetic circuits within such large space has remained absent.

Here, we introduce `BiosInt`, an allosteric transcription factor-based genetic biocircuit, with quasi-integral adaptation control capabilities that can be used to increase robustness and performance of biomanufacturing engineered strains. To test the capabilities of the circuit, we have analysed its performance for the full set of metabolic pathway topologies found in the bio-based chemical production space of compounds with industrial interest. For a given implementation of the `BiosInt` circuit, we carried out a multiobjective optimization process that provides the inverse control solution of optimal topology starting from any given pathway configuration and enzyme expression ratios. Next, synthetic datasets are generated to train machine learning-based predictive models with the data obtained from the simulations of each topology by varying enzyme expression levels and their corresponding concentrations. We validated the models by showing their ability to predict the best control topology for a given set of enzyme ratios.

As a proof-of-concept, we show its application to the design of the genetic constructs expressing flavonoid production pathways, providing optimal performance for the `BiosInt`-mediated dynamic regulation. This circuit inverse design and its application to dynamic control in biomanufacturing paves the way for a future design pipeline in biofoundries delivering more robust and efficient sustainable bioproduction processes.

## 1 Introduction

Microbial cell factories are increasingly been used in bio-based production processes as green alternatives to chemical processes based on fossil resources. In order to make bioproduction processes environmentally sustainable and technoeconomically viable, it is required that they operate efficiently in a circular manner transforming organic feedstocks such as industry and households waste into valuable consumer end products [1]. However, industrial-scale production of bio-based chemicals in the circular green bioeconomy still faces inefficiencies arising from scaling up challenges. To overcome present biomanufacturing inefficiencies, biofoundries have been established as automated facilities where microbial cell factories operate with high efficiency by means of the systematic application of techniques from synthetic biology, automation, and artificial intelligence [2, 3]. Similarly, another cornerstone of efficient microbial biomanufacturing is the introduction of dynamic regulation in production pathways [4]. A common agreement arising from such strategies is that regulation of synthetic metabolic pathways is becoming necessary in modern manufacturing because production strains might be generally prototyped in controlled laboratory conditions [5], but the processes would eventually scaled-up into industrial fermentation conditions, where cell factories will be subject to harsh conditions eliciting stress responses, negatively impacting efficiency [6]. Dynamic control of the synthetic metabolic pathways [7, 8], thus, provides regulation strategies mitigating the loss in efficiency through active control of pathway expression. Yet, a systematic approach for selecting and implementing genetic circuits within such large space has remained largely absent. Biosensor-mediated control of metabolic pathways has been proposed as a strategy to improve the performance of those bioprocesses. By linking industrial biosynthetic pathways to measurable metabolites through biochemical transformations, a highly interconnected regulatory design space is unveiled, offering new opportunities for pathway dynamic control.

Several strategies for dynamic control have been proposed in metabolic engineering and synthetic biology [9, 10]. They are generally based on the introduction of a biosensing system [11], followed by a genetic control circuit [12, 13]. In a similar fashion, several design principles were proposed in different studies, often mimicking what could be understood as the basic functionality of the proportional-integral-derivative (PID) controller in a biological context [14], mainly due to the versatility of such basic control strategy, although other controller designs are possible in synthetic biology which could be seen closer to the actual biological feedback actions found in nature [12, 13, 15]. Notably, efforts have been focused on achieving integral control functionality, as it allows error-free or perfect adaptation in biomolecular systems [16]. A well-known early proposal of such funcionality is found in the antithetic feedback motif [17, 18]. In addition to the controller circuit, design can also be focused on the use of multiple biosensors or the selection of the topology of the circuits. For instance, several control topologies have been proposed for the naringenin pathway beyond a single feedback loop, including a cascaded control using multiple biosensors [19] and the use of three control layers [20].

Here, we introduce `BiosInt`, an allosteric transcription factor-based genetic biocircuit, with quasi-integral adaptation control capabilities that can be used to increase robustness and performance of biomanufacturing engineered strains (see Figure 1). To test the capabilities of the circuit, we have analysed its performance for the set of more than 30,000 metabolic pathway topologies found in the bio-based chemical production and detectable space of main compounds with industrial interest. For a given implementation of the `BiosInt` circuit, we carried out a multiobjective optimisation process that provides the inverse control solution of optimal topology starting from any given pathway configuration and enzyme expression ratios. Next, synthetic datasets are generated to train machine learning-based predictive models with the data obtained from the simulations of each topology by varying enzyme expression levels and their corresponding concentrations. We validated the models by showing their ability to predict the best control topology for a given set of nominal enzyme ratios.

**Figure 1.**
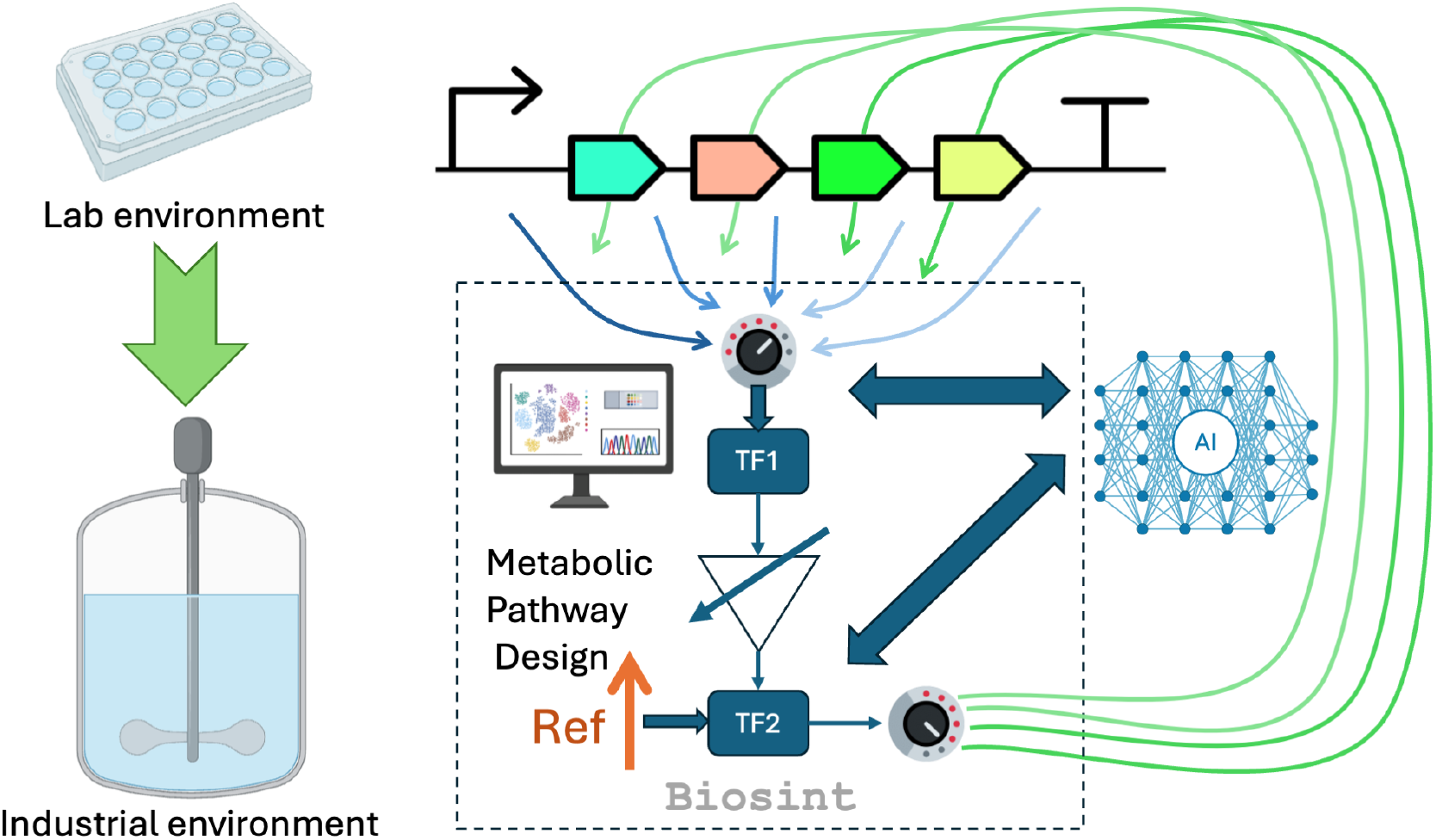
The general design scheme of this study. The `BiosInt` circuit regulates the overexpression of enzymes in the pathway through transcription factor *TF*_2_ based on the feedback measurements obtained from transcription factor *TF*_1_, modulated by the reference *Ref*. An AI-based predictive system selects the optimal topology for sensing and regulation depending on the initial metabolic pathway design. In that way, strains developed in a controlled lab environment become more robust and can overcome stress in industrial production environments.

As a proof-of-concept, we show its application to the design of the genetic constructs expressing flavonoid production pathways, providing optimal performance for the `BiosInt`-mediated dynamic regulation. By introducing this novel circuit inverse design and its application to dynamic control in biomanufacturing, we pave the way for a future design pipeline in biofoundries delivering more robust and efficient sustainable bioproduction processes.

## 2 Materials and methods

### 2.1 Data sets

#### DetSpace sources

The list of 484 compounds of industrial interest was obtained from a curated map of bio-based chemical compounds [21]. This list was reviewed and expanded accordingly. The 436 detectable molecules were sourced from Sensbio, a comprehensive database of biosensors [22]. These databases were combined and cross-linked to obtain the reference metabolic pathways on the DetSpace web server, through the use of reaction rules [23]. These reaction rules are able to connect detectable and producible compounds through the use of bioretrosynthesis.

#### Kinetic constants

The kinetic constants of the enzymes were obtained from the Brenda database [24] in the case of specific pathways, like naringenin. For generic enzymes, a range of values between 10^*−*2^ and 10^5^ was used.

### 2.2 Determination of topologies

Starting with the compound of industrial interest, the metabolic pathway was traced backwards bioretrosynthetically through the iterative application of reaction rules until the precursor compounds were found to be present in the host organism, *Escherichia coli*. With this information, a graph representing the pathway was created, removing compounds that can be considered trivial, such as water or other cofactors, to obtain the Weisfeiler Lehman graph descriptor hash [25]. This method ensures identical results for isomorphic graphs and different results for non-isomorphic, allowing us to classify the metabolic pathways according to whether they are isomorphic or not. The principal component decomposition (PCA) of the graph set based on the topology descriptor started from 43 components, and they were reduced to 23 maintaining the cumulative explained variance ratio above 0.95. In order to classify the set, the topologies were codified as shown in Figure 2. For the bioproduction design space, 48 different topologies were found; and nearly 35,000 for the biodetectable space.

**Figure 2.**
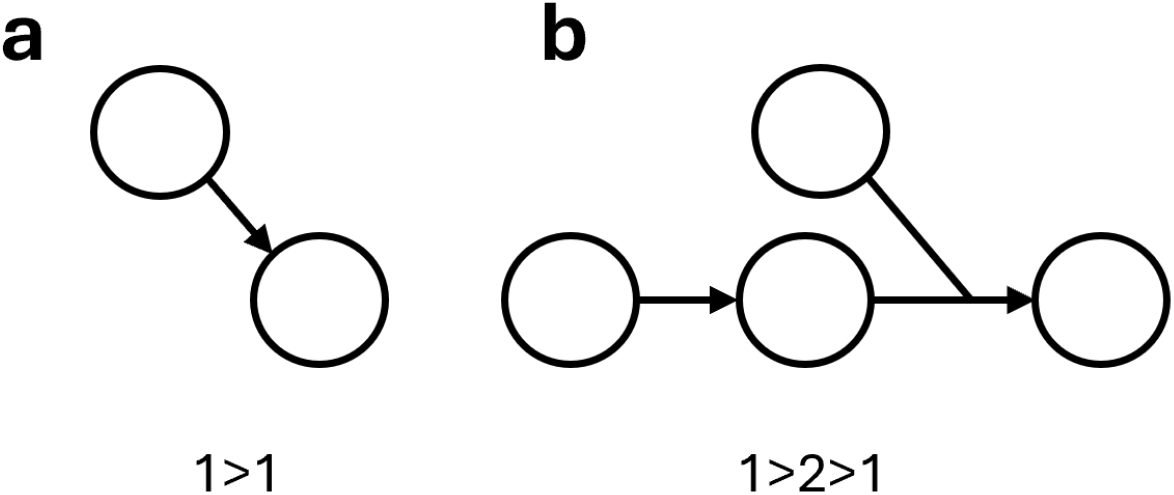
Diagram of two topologies present in DetSpace and their corresponding coding. (a) This topology presents a single reaction with one substrate and one product, therefore, the graph is coded as 1*>*1. (b) This topology has two reactions, the second one with two substrates, therefore, the graph is coded as 1*>*2*>*1.

### 2.3 Modeling the pathway response

Reactions in the pathway follow the Michaelis-Menten equation 1, where [*Product*] and [*Substrate*] are the concentrations of each compound. [*E*] represents the enzyme concentration and *k*_*cat*_ and *K*_*m*_ its kinetic constants. *K*_*d*_ is the degradation constant of the product:

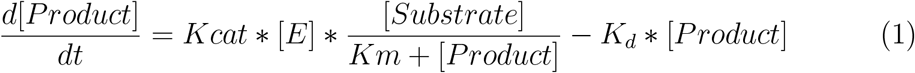

When constructing the generic cases, random values within the range [10^*−*2^, 10^5^] mM/s were assigned to the kinetic constants *k*_*cat*_ and *K*_*m*_ for each of the enzymes in the pathway. A constant value was set for the concentrations of the input compounds, with a degradation constant of 0.01 mM/s. The enzyme concentrations are within the range [0.01, 100] mM. For multiple substrates, we used a generalized Michaelis-Menten equation.

### 2.4 Modeling the BiosInt dynamic behavior

Based on the previously modelled route, a control was implemented on the dynamic response. The steady state values for each metabolite concentration were used as the initial conditions. A designated implementation of the `BiosInt` parameters was defined and we assumed that such configuration is the one available for the controller device in all experiments. The controller operation requires two user-defined parameters that determine the control topology: the enzyme, whose expression is being regulated by the controller and the metabolite that will be detected by using an available biosensor.

The simulation of the genetic circuit consists of two stages. First, the regulated is simulated until it returns to the steady state. In the second stage, a perturbation is introduced into one of the pathway precursor metabolites by lowering its concentration, and the simulation is performed until the system reaches the new equilibrium state. The model equations are as follows:

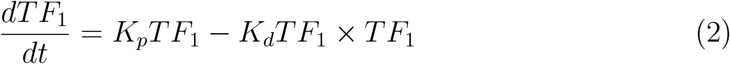

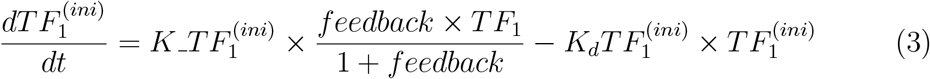

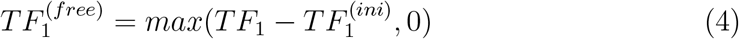

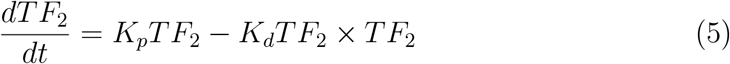

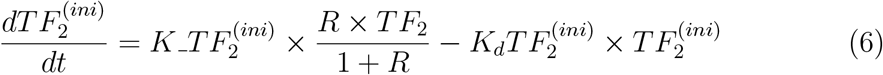

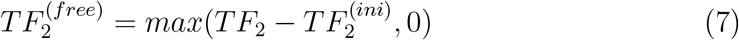

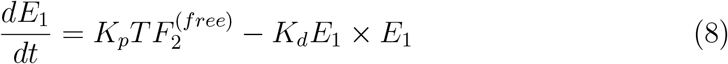

The definitions of each parameter and the value assigned are shown on Table 1.

**Table 1:** Parameters of the BiosInt model.

| Table 1: Parameters of the BiosInt model. |  |  |
| --- | --- | --- |
| Parameter | Definition | Value |
| $K_p TF_1$ | Promoter *Pro1* strength | 1 (mM/s) |
| $K_p TF_2$ | Promoter *Pro2* strength | 100 (mM/s) |
| $K_p E_1$ | Promoter *Pro3* strength | 1e-3 (mM/s) |
| $K\_TF_1^{(ini)}$ | Constant for production of $K\_TF_1^{(ini)}$ | 0.5 (mM/s) |
| $K\_TF_2^{(ini)}$ | Constant for production of $K\_TF_2^{(ini)}$ | 0.5 (mM/s) |
| $K_d TF_1$ | Degradation rate of $TF_1$ | 0.1 (mM/s) |
| $K_d TF_2$ | Degradation rate of $TF_2$ | 0.1 (mM/s) |
| $K_d TF_1^{(ini)}$ | Degradation rate of $TF_1^{(ini)}$ | 1 (mM/s) |
| $K_d TF_2^{(ini)}$ | Degradation rate of $TF_2^{(ini)}$ | 1 (mM/s) |
| $K_d E_1$ | Degradation rate of $E_1$ | 1 (mM/s) |
| $TF_1$ | Concentration of $TF_1$ | Variable |
| $TF_2$ | Concentration of $TF_2$ | Variable |
| $TF_1^{(ini)}$ | Concentration of $TF_1^{(ini)}$ | Variable |
| $TF_2^{(ini)}$ | Concentration of $TF_2^{(ini)}$ | Variable |
| feedback | Concentration of the biosensor | Variable |
| $R$ | Concentration of the reference species R | Variable |

### 2.5 Selection of the optimal topology

The control topology is defined by two selectable parameters: the reaction being controlled and the metabolite used as a biosensor. Figure 3 shows two examples of possible control topologies for the 1*>*1*>*1 topology. In these examples, the metabolite used as a biosensor, *B* or *C*, and the reaction being controlled, *R1* or *R2*, change. The graph shows how by modifying this configuration we obtain different dynamic response for the final product *C*.

**Figure 3.**
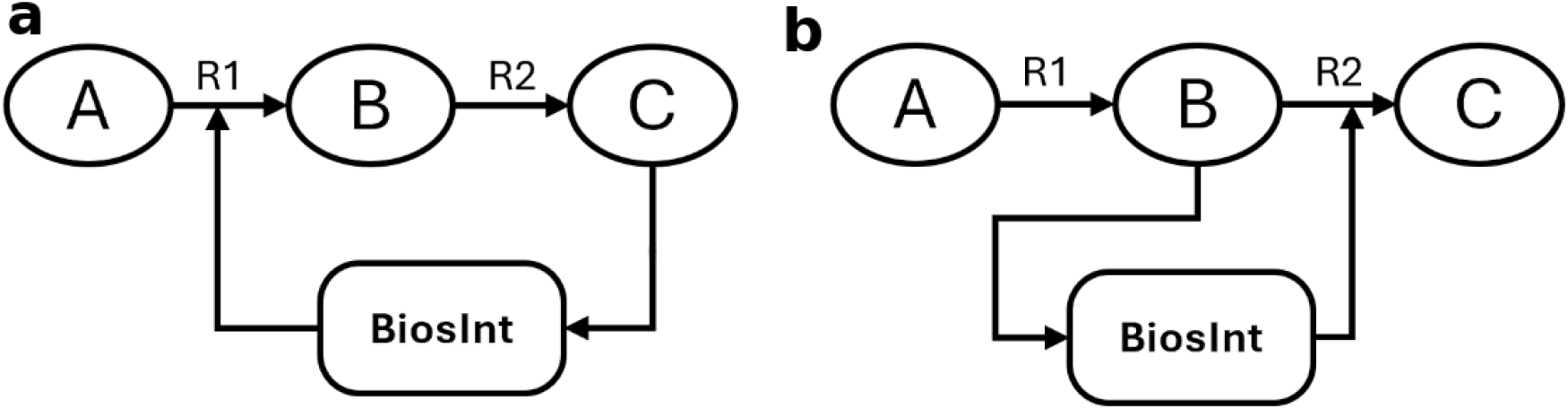
Two examples of control topologies for 1*>*1*>*1. (a) Topology 1 (top figure) uses *C* as biosensor and regulates expression of enzyme *R*_1_. (b) Topology 2 (bottom figure) uses *B* as biosensor and regulates *R*_2_.

For each of the different pathway topologies, pathway responses were calculated by varying the two control parameters using the models described above. Four values were obtained in each simulation: *O*_1*N*_, *O*_1*P*_, *O*_2*N*_, *O*_2*P*_, which are defined as follows: *O*_1_ corresponds to the concentration of the final product of the pathway; *O*_2_ is the average concentration of the enzymes; *N* and *P* indicate the two stages of the simulation, with *N* being the moment prior the perturbation and *P* being the final moment of the simulation. With these values, a score was assigned to each combination of control parameters as: *O*_1*P*_ /*O*_2*P*_.

For each pathway topology, this process was carried out for different enzyme concentrations in order to determine the optimal control topology at different context conditions.

### 2.6 Predictive model

A predictive model based on a feed-forward neural network was constructed from the data previously obtained. For a given topology, the training set consisted of different combinations of enzyme concentrations as inputs, and the outputs were the optimal control topologies, encoded through one-hot-encoding. For each enzyme, 9 concentration values were defined within the range [0.01, 100] and all possible combinations between the different enzymes in the pathway were explored. In the case of the topologies 1*>*1 and 2*>*1, more data need to be generated as there is only one reaction, so 100 values within the range were used. The dataset was divided into 80% training and 20% test. `Tensorflow` and `Keras` were used to train the neural network. Accuracy was used as a measure of performance. For each topology, cross-validation was repeated 100 times, and the obtained mean accuracy was used as a result. The neural network has a 4-layer architecture. The intermediate layers consist of a layer of 64 nodes with sigmoid activation function and a dropout layer. The output layer has a softmax activation function. The optimizer is an Adam optimizer and the loss function is categorical cross entropy from `Keras`.

### 2.7 Use case: dynamic regulation of the naringenin pathway

The described general workflow was also applied in a focused study for the naringenin pathway, see Figure 4. The pathway consists of 4 reactions, with the third one using two products (topology code 1*>*1*>*2*>*1). *k*_*cat*_ and *K*_*m*_ values for each reaction were obtained from the Brenda database [24]. The compounds involved in the pathway, except for naringenin chalcone, can be associated with an allosteric transcription factor, and, therefore, they can be detected by a biosensor for dynamic regulation. In total, thus, there are 20 possible optimal topologies, i.e., *(4 reactions* × *5 biosensors)*. As we considered 9 possible values for each enzyme concentration, the resulting dataset contains 6,500 different combinations, which can be then used to train the predictive model, in a similar fashion as for other topologies considered in the study.

**Figure 4.**
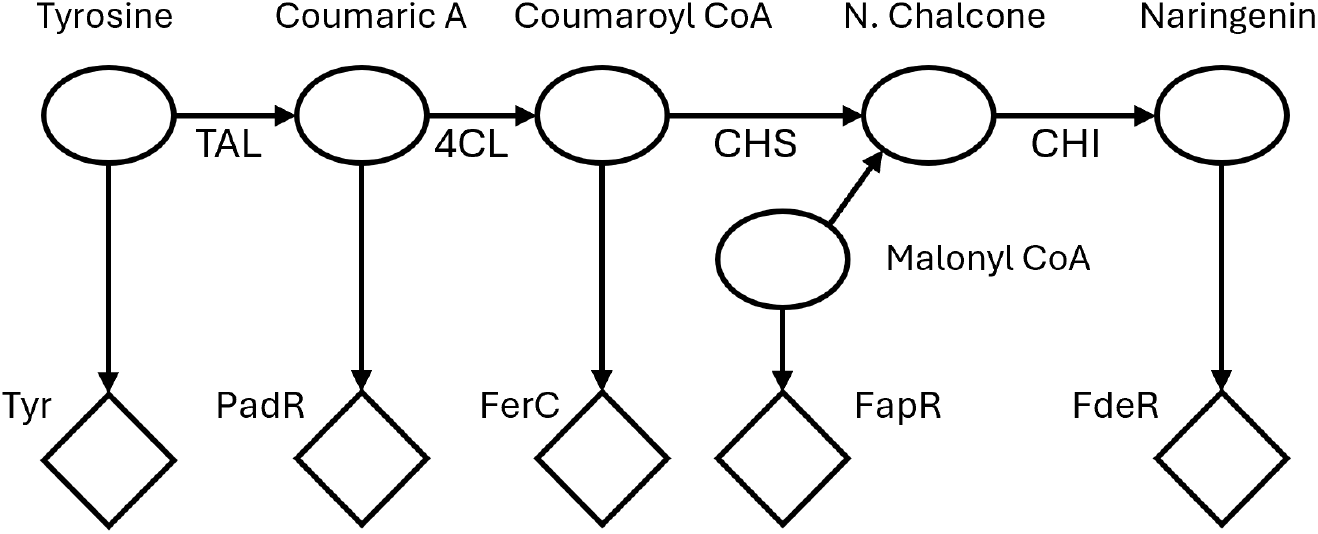
Production pathway of the flavonoid naringenin, showing the enzymes involved in each reaction (TAL, 4CL, CHS, CHI), as well as the allosteric transcription factors activated by metabolites in the pathway (Tyr, PadR, FerC, FapR, FdeR).

## 3 Results and Discussion

### 3.1 Determination of the topological distribution of the bioproduction and biodetectable design spaces

Synthetic bioproduction pathways are established by a set of enzymes that determine a metabolic route for the conversion of endogenous metabolites into the desired target products. Several studies have determined a consensus reference set of approximately 500 pathways that contains the most industrially-relevant bio-based targets for the bioeconomy [21]. Those pathways contain a large diversity in terms of their topology content, meaning by this that even though some of the pathways are linear chains of reactions, others contain branched topologies where at least one of the reaction steps depend on two or more reactants. Branched steps need all their reactants to be produced in order for the reaction to proceed, therefore creating more complex pathway topologies. Considering *Escherichia coli* as the chassis organism, we found 48 distinct topologies contained in the aforementioned reference industrial pathway set. Figure 5 shows their topology distribution, which approximately follows an exponential law. Most of the pathways had a simple linear distribution containing a few number of enzymes. However, branched distributions were also common, especially for low-length pathways.

**Figure 5.**
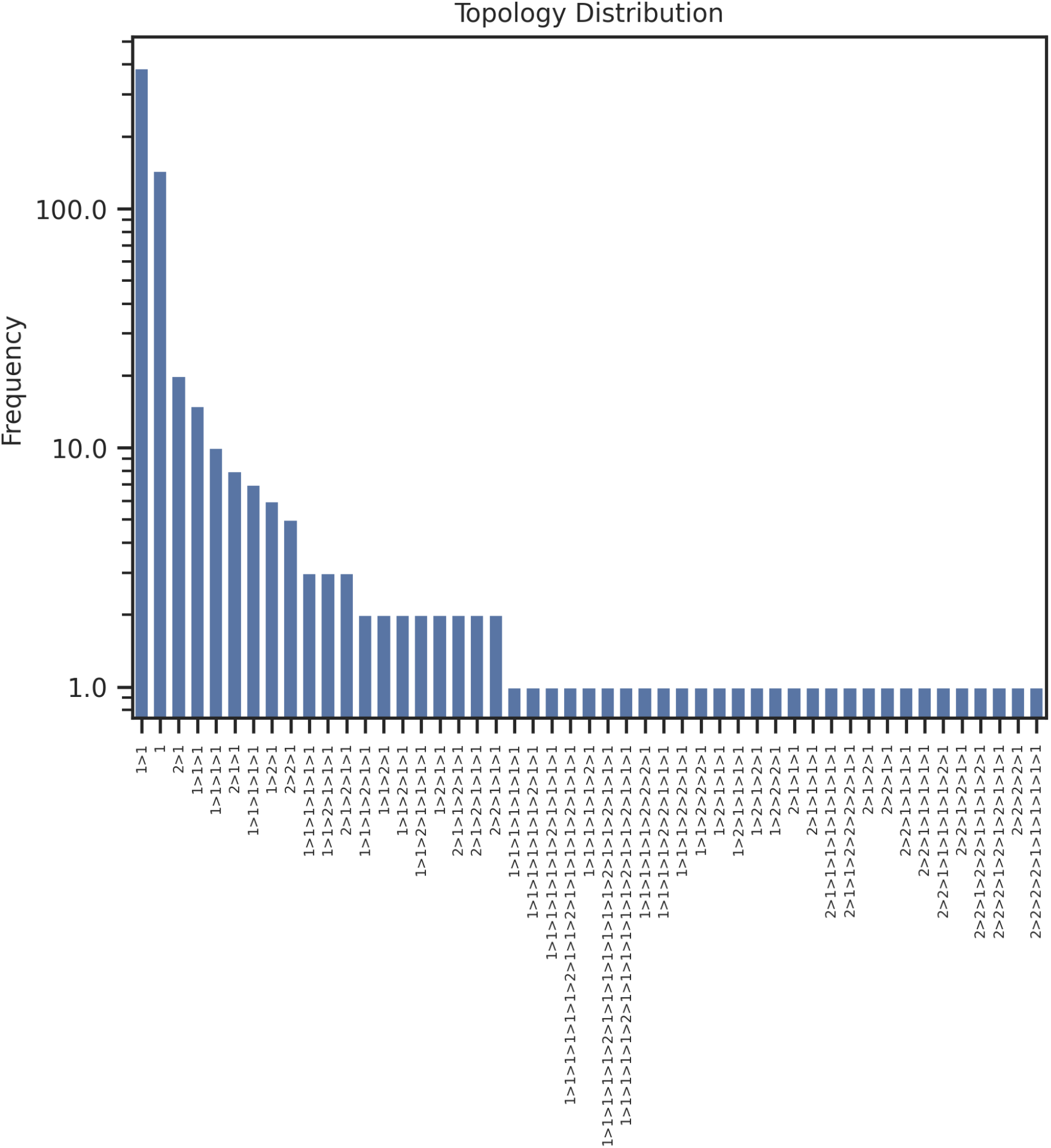
Distribution of topologies in the top 500 industrially-relevant bio-based targets pathways, expressed in *E. coli* as chassis organism. Enzymes in the pathway topology are represented by the ‘*>*’ symbol, and are preceded by their number of precursors. Branched steps occur when the number of precursors is greater than one.

Our purpose is to evaluate the viability of establishing and automating the design of a dynamic regulation strategy for each of those topologies through the `BiosInt` circuit, as its behavior that is close to an integral controller should facilitate the development of producing strains with an increased robustness. In order to implement such strategy, we need first to determine those metabolites and intermediates in the pathway that can be sensed through an allosteric transcription factor, and therefore candidates to establishing a feedback loop in the pathway through a regulation circuit that actuates into the expression of one of the genes associated with the enzymes in the pathway.

In a previous study, we introduced the tool DetSpace [23], which provides the full set of metabolites in the bioproduction design space that can be detected, directly or through a sensing pathway by means of an allosteric transcription factor. Therefore, we can calculate by means of the DetSpace tool the topological distribution of the detectable design space, i.e., a metabolic circuit consisting of a production pathway and a detection pathway connecting the target metabolite to an inducible transcription factor. In order to analyze the topological distribution of the collection of circuits for the biosensing pathways under study, a PCA decomposition was performed on the topology descriptor (see Materials and Methods). On the graph were represented the two most important. The resulting distribution is shown in Figure 6 for the two main components, highlighting the exponential distribution of the number of occurrences of each type of topology.

**Figure 6.**
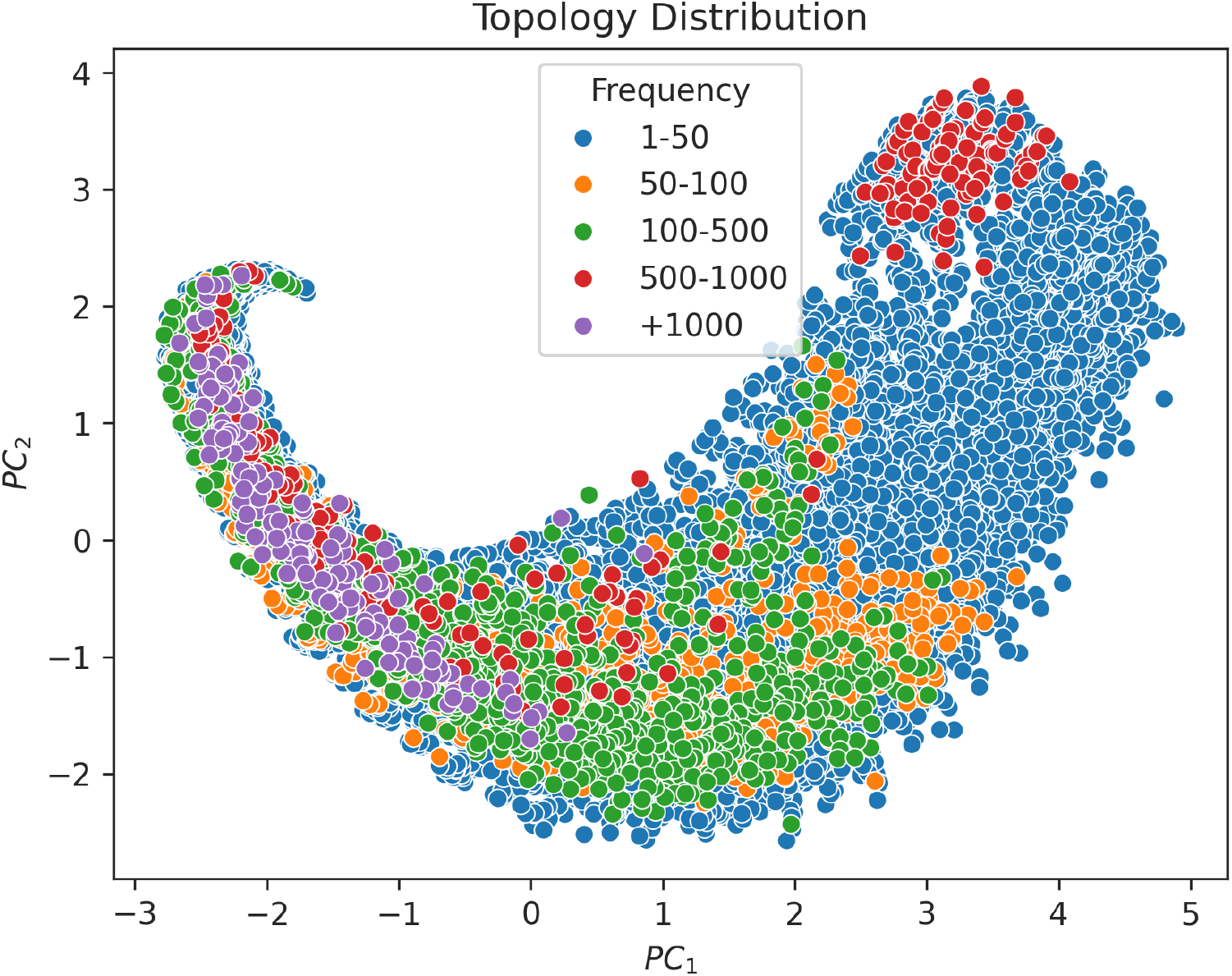
Distribution of topologies present on DetSpace across the two main components *PC*_1_, *PC*_2_ after a PCA was applied to their descriptor vector. The colors represent the frequency with which each topology appears.

In the obtained distribution, some clusters were observed with shared features, suggesting an underlying pattern. For instance, a cluster composed of topologies with a frequency between 500 and 1000 was formed (top right corner in Figure 6) whose topologies turned out to be those with the longest average length. Similarly, a cluster with frequencies greater than 1000 is also found on the left side of the figure, where the topologies contained in the cluster are those with the highest presence of bifurcations, highlighting the complexity of the set. In both cases, topologies always presented bifurcations at the initial reactions. For the rest of the distribution groups, no notable clusters are apparent. Unlike in bioproduction, all linear topologies have low frequency, between 1 and 50. This may be explained because DetSpace design space has been expanded to contain as many routes as possible, generating more complex topologies.

### 3.2 Introducing the BiosInt dynamic regulation circuit

Our goal is to develop an allosteric transcription factor-based controller circuit that can implement the dynamic regulation of biosynthetic pathways based on the optimal combination between biosensors and transcriptional actuators of the genes inolved in the pathway. To that end, Figure 7 shows the basic configuration of the `BiosInt` circuit. The circuit consists of two allosteric transcription factors that work as repressor-activator. Transcription factor *TF*_1_ works as a biosensor for the sensed molecule *T*, which inhibits the biosensor. The transcription factor activates the promoter *Pro*_2_ that expresses allosteric transcription factor *TF*_2_, which is continuously inhibited by effector *R* that provides a continuous reference level and can be either produced by some enzyme *REF*_1_ or fed into the medium. Non-inhibited *TF*_2_ will activate a promoter *Pro*_3_ that, in turn, allows overexpression of a production pathway enzyme *E*_1_, which in the example represented in Figure 7 is the one that directly overproduces the sensed target *T*.

**Figure 7.**
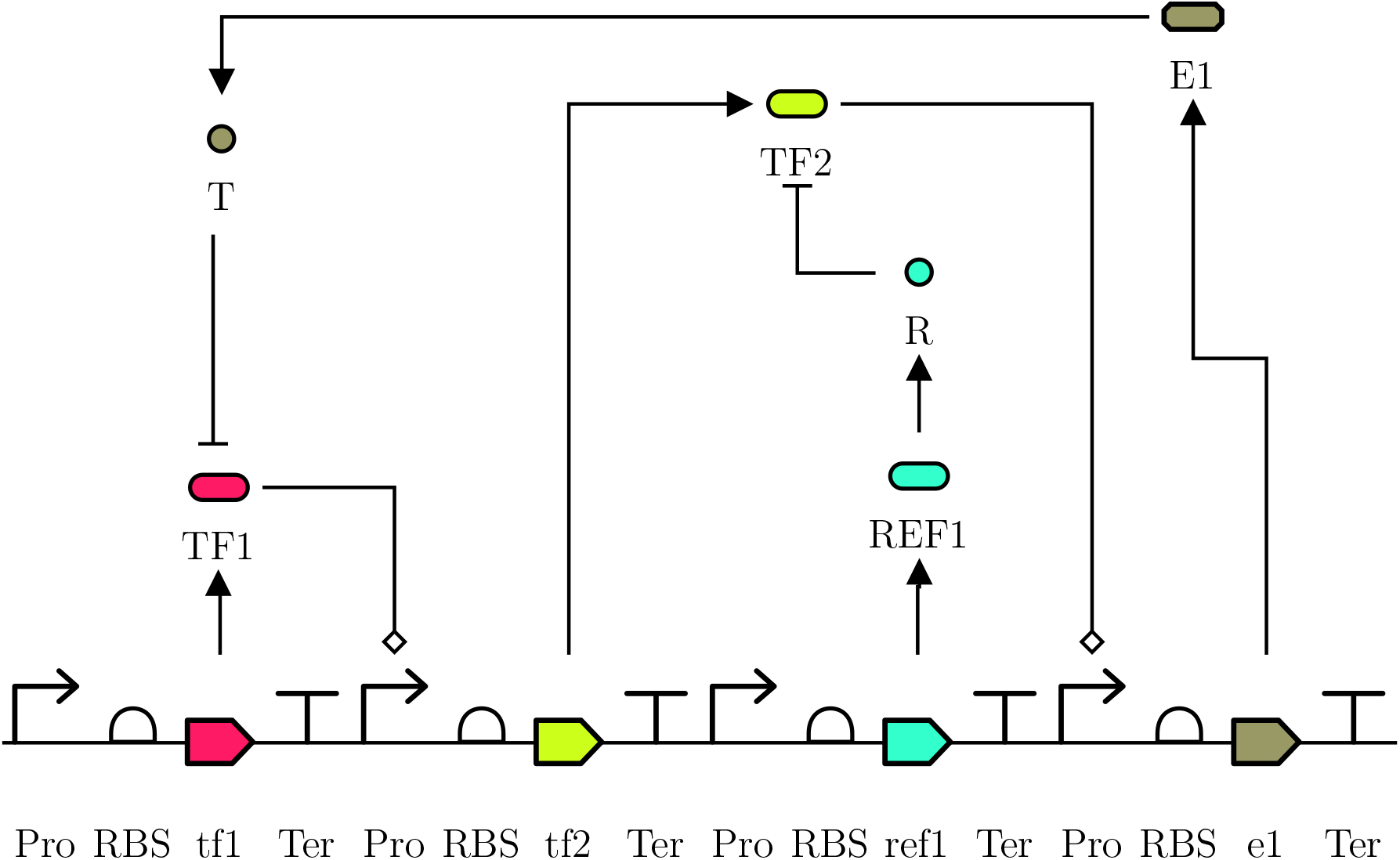
The basic configuration of the `BiosInt` circuit regulates the expression of enzyme *E* through a cascade interconnection of transcription factors *TF*_1_ and *TF*_2_, which are, in turn, allosterically inhibited by *T* and *R*, respectively.

In case that the intracellular availability of target *T* decreases, *TF*_1_ will increase the expression of *TF*_2_ and therefore, there will be more a higher availability of non-inhibited triggering the overexpression of enzyme *E*_1_ and therefore more target compound *T* could potentially be produced, provided that the theoretical maximum titer has not been yet reached. This strategy, thus, should be able to correct deviations from the initial production titers due to changes in the medium conditions, as long as the system is kept away from the saturation regime.

Interestingly, the `BiosInt` circuit could be eventually tuned to a desired regulation action by changing the levels of reference species *R* present in the media, allowing therefore an external control of the system. Figure 8a shows different dynamic responses for a generic pathway with topology 1*>*1, according to the level of reference species *R*. As mentioned above, a higher value for the reference results in a lower production of metabolite *T*. Figure 8b shows the dose-response curve of the target for different values of the reference *R*. In this example, the reference *R* allows the fine-tuning of the product end-value within a range given approximately by 14-17 mM.

**Figure 8.**
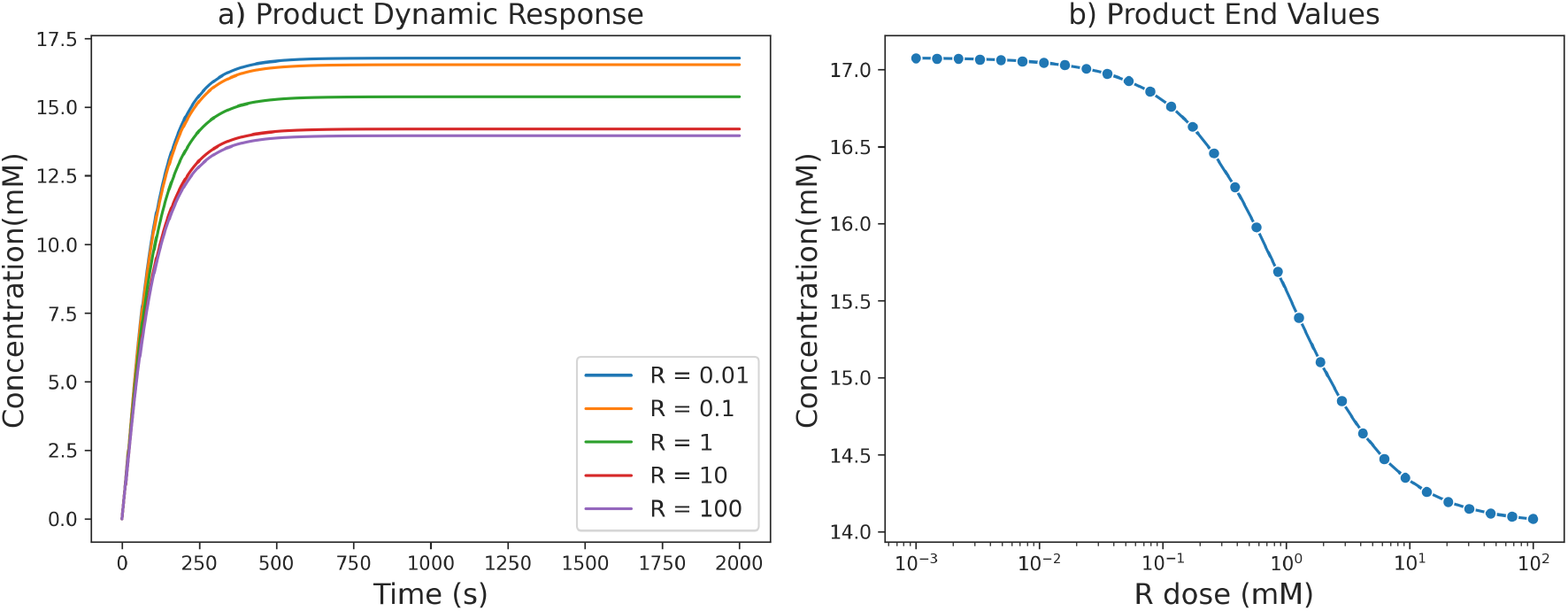
Dynamic responses of a pathway with topology 1*>*1, according to the level of reference species *R*. a) Product dynamic response; b) Product end values dose-response curve depending on reference *R*.

### 3.3 Mapping optimal topologies depending on pathway context

Once a target compound has been selected, there is a limited selection of gene variants for each enzymatic step in the production pathway. Such limited selection is therefore a design constraint, meaning that only a reduced set of possible choices is available. Each choice of gene variant, and associated regulatory elements like promoters and RBSs will imply a different balance between the expression levels of the enzymes involved in the pathway. We investigated how this design choice might determine the optimal control topology for the `BiosInt` closed-loop regulation. The control topology is defined by two parameters: the reaction being controlled and the metabolite detected by the biosensor. The optimal topology is defined as the topology that provides the best control, according to some criteria, for a specific pathway in a particular context, i.e., for the enzymes selected in the pathway. In the study, the best control was defined as the one that achieves the best trade-off between the highest concentration of the final product after a perturbation and the associated metabolic burden (see Materials and Methods).

As shown in Figure 9, the optimal regulation topology depends on the context for each pathway class. In this example, a two-enzyme pathway (*E*_1_, *E*_2_) was chosen. Depending on the variants, steady-state concentrations of the enzymes in the pathway will vary. We explored how this design choice could impact the selection of the best feedback topology, namely, the selection of the regulated enzyme and feedback metabolite. As expected, low concentrations of *E*_1_ or *E*_2_ are usually compensated by the over-expression of the depleted enzyme and feedback of the enzyme precursor. Interestingly, when concentrations become closer for both enzymes, there is a transition region where the optimal topology is transferred to another metabolite in the pathway. As concentrations become split apart, the optimal topology converges to the enzyme present at lowest concentration.

**Figure 9.**
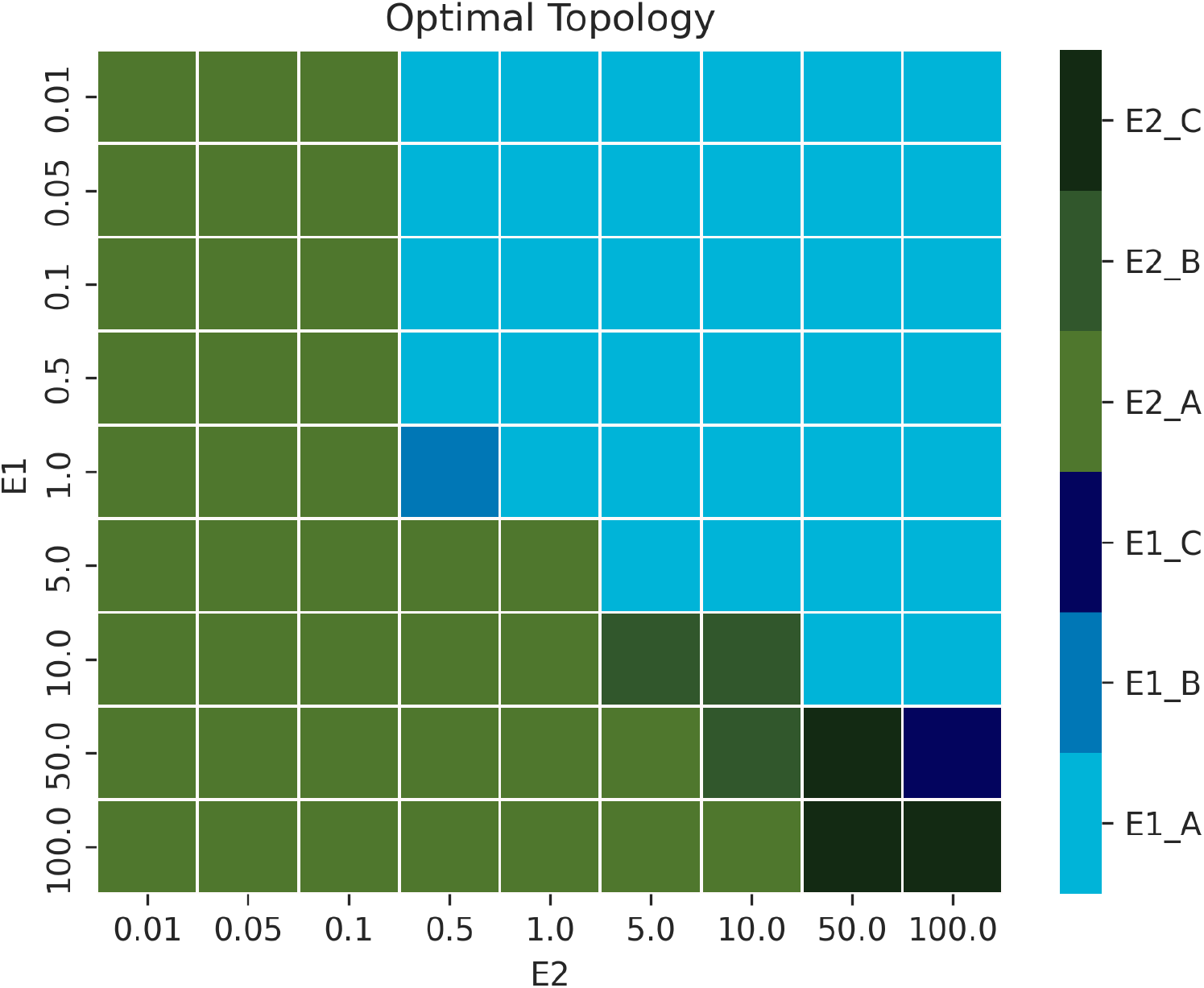
Optimal topology depending on enzyme concentration. In this example pathway, there are two enzymes *E*_1_ and *E*_2_, and three metabolites *A, B*, and *C*. Depending on the design context, steady-state concentrations of the pathways might vary. The optimal feedback topology for `BiosInt`-based regulation is shown in the heatmap, where each color code represents a pair regulated enzyme-feedback metabolite.

**Figure 10.**
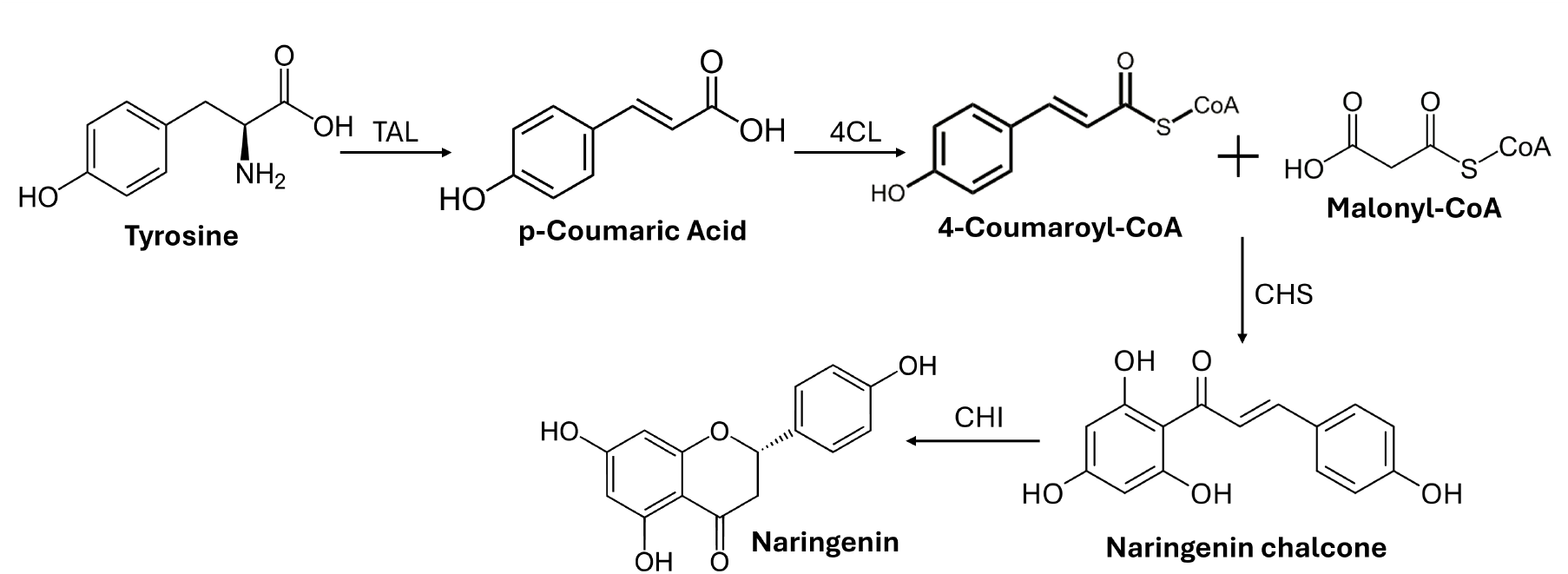
The naringenin bioproduction pathway and the associated intermediate metabolites.

Therefore, there is a transition region where we observe that the problem of choosing the optimal topology has no straightforward solution and shows a complex dependency with respect of the initial enzyme concentrations. Interestingly, the situation becomes more complex as the complexity of the pathway increases, either in length or in the presence of branched steps, as shown in the topology distribution in Figure 5.

### 3.4 Prediction of the optimal topology

Results from previous section suggest that as the complexity of the pathway increases, the selection of the optimal topology for a given implementation of the `BiosInt` circuit becomes more challenging. This predictive inverse optimal topology design problem can be stated as follows:

#### Given

1. A target metabolite and a chassis organism;
2. An implementation of a metabolic pathway design, carried out through metabolic engineering and synthetic biology methods;
3. A `BiosInt` controller design, tuned to a designated regulation function;

#### Find

An optimal topology in terms of regulated enzyme and feedback metabolite according to the desired criteria for pathway performance (production, robustness, efficiency, etc.).

In order to solve this problem, we introduce here a machine-learning based predictive system. The system should be able to learn an optimal solution to the problem based on the given context and the defined biomanufacturing objectives, which in this case were defined as maximizing the ratio between recovery after a perturbation in the production of the target compounds with respect to the metabolic burden elicited by over-expressing the regulated enzyme. To that end, we generated a training set consisting of the synthetic data obtained from the simulation for each topology and possible combinations of initial pathway configurations, defined by the enzyme concentrations, and the performance indicators obtained by the steady-state values of the regulated pathways (see Materials and Methods). The training set was then use to build and validate a predictive model capable of predicting the optimal topology based on the context.

Several machine learning approaches were investigated. In order to assess the complexity of the problem under study, we compared the performance of a Bayesian predictor trained with the generated data, with that of a neural network. Notably, results obtained by the neural network offered a clear improvement with respect to the Bayesian predictor. As shown in Table 2, we obtained the average performance, based on prediction accuracy in the cross-validation, and its associated standard deviation for each of the topologies present in the training set, both for the neural network and the Bayesian. It can be seen that in all cases the neural network outperformed the Bayesian model, indicating as expected that the problem requires of a powerful training system to be able to solve accurately the topology selection problem. In general, the increase in performance was more apparent for those pathways with higher length and number of branches, with cases with an increase in performance of 20%, like in the case of the 1*>*2*>*2*>*2*>*1 topology.

**Table 2:** Performance comparison between neural network model (NN) and Bayesian predictor (Bayesian), according to rhe cross-validation mean accuracy.

| Topology | Mean (NN) | Mean (Bayesian) | Std (NN) | Std (Bayesian) |
| --- | --- | --- | --- | --- |
| 1>1 | 0.9920 | 0.9793 | 0.0020 | 0.0103 |
| 1>1>1 | 0.8864 | 0.8509 | 0.465 | 0.1012 |
| 1>1>1>1 | 0.9013 | 0.7936 | 0.0087 | 0.0338 |
| 1>1>1>1>1 | 0.9314 | 0.7655 | 0.0030 | 0.0299 |
| 2>1 | 0.9929 | 0.9732 | 0.0020 | 0.0119 |
| 1>2>1 | 0.9085 | 0.8958 | 0.0191 | 0.0684 |
| 2>1>1 | 0.8866 | 0.8634 | 0.0233 | 0.0891 |
| 2>2>1 | 0.9119 | 0.8834 | 0.0209 | 0.0812 |
| 1>1>2>1 | 0.9134 | 0.8613 | 0.0059 | 0.0295 |
| 1>2>1>1 | 0.9384 | 0.8616 | 0.0057 | 0.0334 |
| 2>1>1>1 | 0.9365 | 0.8956 | 0.0060 | 0.0263 |
| 2>2>1>1 | 0.9448 | 0.8924 | 0.0169 | 0.0272 |
| 2>1>2>1 | 0.9256 | 0.8504 | 0.0177 | 0.0266 |
| 1>1>2>1>1 | 0.9423 | 0.7071 | 0.0069 | 0.0206 |
| 1>2>1>1>1 | 0.9588 | 0.8104 | 0.0065 | 0.0142 |
| 2>1>1>1>1 | 0.9449 | 0.7537 | 0.0061 | 0.0179 |
| 2>2>1>1>1 | 0.9400 | 0.7953 | 0.0068 | 0.0133 |
| 2>2>2>2>1 | 0.9487 | 0.7504 | 0.0066 | 0.0219 |
| 1>2>2>2>1 | 0.9502 | 0.7461 | 0.0056 | 0.0198 |

### 3.5 The case of the naringenin pathway

The production pathway for the naringenin flavonoid offers an interesting case study, since most of the metabolites involved in the pathway can be detected by a transcription factor and, therefore, they can serve as the feedback point for the `BiosInt` circuit. As it has been said before, naringenin chalcone is the only one that does not have an allosteric transcriptor factor associated. For this specific study, we have added a variation. In order to produce a perturbation in the compound production, we will study the case in which the perturbation appears as the depletion of one of the compounds involved in the pathway bifurcations.

The obtained optimal topologies in the combinatorial design space of initial enzyme concentrations showed that regulation of the CHS and CHI can provide the best control, as they appear in approximately 50% of the combinations. This result may be related to the fact that these two reactions are the ones causing a bottleneck in the pathway. Therefore, it is important actuating on them to alleviate the bottleneck effect. On the other hand, the frequency with which each metabolite appears as a sensor is shown in Table 3. It can be seen that the compound acting as the best sensor is, in most cases, the one that undergoes the perturbation. According to this observation, it might be reasonable to think that detecting the perturbation directly produces the best results in control compared with other possible indirect measurements. In the case of a tyrosine deficiency, a significant percentage of cases were also found where the second branched compound, malonyl-CoA, present downstream in the pathway with respect to tyrosine, was the optimal feedback compound, which might be due to cases where there is a high availability of the tyrosine-consuming enzyme TAL, and therefore feedback in the dynamic regulation is shifted towards malonyl-CoA.

**Table 3:** Ratio of occurrence of each of the compounds in the naringenin pathway acting as biosensors in the optimal topology within the combinatorial design space.

| Metabolite | Occurrence rate<br>for tyrosine deficiency | Occurrence rate<br>for malonyl-CoA deficiency |
| --- | --- | --- |
| Tyrosine | 74.46 % | 0.89 % |
| Coumarate | 3.12 % | 1.48 % |
| Coumaroyl-CoA | 0.93 % | 1.17 % |
| Naringenin | 0.00 % | 2.33 % |
| Malonyl-CoA | 21.49 % | 94.13 % |

The combinatorial design space of the naringenin producing pathway, modeled by their steady-state enzyme concentrations was simulated in order to generate a training set to build and cross-validate a predictive model for optimal dynamic regulation topology selection. As in the previous case, the neural network offered a better performance than the Bayesian model, as shown on Table 4. For the naringenin case, the obtained increase in performance was in excess of 20%, achieving an accuracy close to 95% in the neural network.

**Table 4:** Performance comparison between neural network model (NN) and Bayesian predictor (Bayesian) for the naringenin pathway.

|  | NN | Bayesian |
| --- | --- | --- |
| Mean | 0.9454 | 0.7042 |
| Std | 0.0078 | 0.0128 |

## 4 Conclusions

In summary, we have introduced here a regulation circuit, `BiosInt` and its associated AI-based design tool `biosint.ai`, with the ability of selecting the optimal topology for increasing strain robustness given an initial metabolic pathway design. As the demand for industrial biomanufacturing processes increases due to the urging need of transitioning towards a global sustainable bioeconomy, we anticipate a growing number of bio-based production processes having the need of this type of regulation strategies to confer robustness to the system in order to make the overall process viable and sustainable.

Our approach differs from other proposed solutions, which depend on the availability of a detailed knowledge of the pathway mechanistic dynamics and assume that the regulation circuit can be easily tuned in function of the chosen design solution. On the contrary, in this study we assume that the regulation circuit, named `BiosInt` is already built and its functionality has been experimentally characterised and, in addition, the desired pathway has been already designed based on metabolic engineering considerations and well-established methods. Therefore, the main goal of our approach is to develop a predictive system that can make an automated decision system that can select the best regulation topology to choose from within the large combinatorial set, trained on the previously obtained data, rather than having a focus on a particular class of pathways or compounds. We believe that this compound-agnostic approach that solves the inverse design problem based on the availability of pathway biosensor capabilities, will facilitate the adoption of the proposed dynamic regulation framework within a general Design-Build-Test-Learn (DBTL) workflow by easing the overall design process, boosting in that way progress towards advanced next-generation industrial biomanufacturing.

## Acknowledgments

HM, IOM, PC acknowledge support from Generalitat Valenciana through grant CIAICO/2021/159 (SmartBioFab) and MCIN/AEI /10.13039/501100011033 and European Union NextGenerationEU/ PRTR funding through grant TED2021-131049B-I00 (BioEcoDBTL).

The computations were performed on the HPC cluster Garnatxa at Institute for Integrative Systems Biology (I2SysBio), I2SysBio is a mixed research center formed by University of Valencia (UV) and Spanish National Research Council (CSIC), and at the HPC cluster Rigel of the Universitat Politécnica de Valéncia.

## Author contributions

HM performed the simulations, analyzed the data and generated the figures. PC conceived the study and analyzed the data. IOM contributed to the modeling and performance assessment. HM, IOM, PC wrote the manuscript. All authors have given approval to the final version of the manuscript.

## Code availability

The code used to generate the results presented in this study is available at the following GitHub repository: https://github.com/dbtl-synbio/biosint.

